# The Catalytic Core of DEMETER Guides Active DNA Demethylation in Arabidopsis

**DOI:** 10.1101/623843

**Authors:** Changqing Zhang, Yu-Hung Hung, Xiang-Qian Zhang, Dapeng Zhang, Jennifer M. Frost, Fang Liu, Wenyan Xiao, Lakshminarayan M. Iyer, L. Aravind, Jin Hoe Huh, Robert L. Fischer, Tzung-Fu Hsieh

## Abstract

The Arabidopsis DEMETER (DME) DNA glycosylase demethylates the maternal genome in the central cell prior to fertilization, and is essential for seed viability. DME preferentially targets small transposons that flank coding genes, influencing their expression and initiating plant gene imprinting. DME also targets intergenic and heterochromatic regions, and how it is recruited to these differing chromatin landscapes is unknown. The C-terminal DME catalytic core consists of three conserved regions required for catalysis in vitro. We show that the catalytic core of DME guides active demethylation at endogenous targets, rescuing the developmental and genomic hypermethylation phenotypes of DME mutants. However, without the N-terminus, heterochromatin demethylation is significantly impeded, and abundant CG-methylated genic sequences are ectopically demethylated. We used comparative analysis to reveal that the conserved DME N-terminal domains are only present in the flowering plants, whereas the domain architecture of DME-like proteins in non-vascular plants mainly resembles the catalytic core, suggesting that it might represent the ancestral form of the 5mC DNA glycosylase found in all plant lineages. We propose a bipartite model for DME protein action and suggest that the DME N-terminus was acquired late during land plant evolution to improve specificity and facilitate demethylation at heterochromatin targets.

## Introduction

DNA methylation is a covalent modification that influences the transcription of nearby genes and regulates important processes in eukaryotic genomes, including cell differentiation, transposable element (TE) silencing, and genomic imprinting (1, 2). Plant DNA methylation occurs in CG, CHG, and CHH sequence contexts (H = A, C, or T) and is primarily targeted to TEs. Flowering plants and mammals can also exhibit gene body methylation (gbM) in the CG context, generally in constitutively expressed genes, but the function of gbM is not fully understood (3, 4).

Maintaining DNA methylation homeostasis is essential for genome stability, notably in maintaining TE silencing, and for the stable and long-term inheritance of epigenetic information (5, 6). In plants, this is achieved by maintenance and de novo DNA methylation, and by active DNA demethylation (1, 7–10), Active DNA demethylation is catalyzed by a family of novel DNA glycosylases, REPRESSOR OF SILENCING 1 (ROS1), DEMETER (DME), DEMETER-LIKE 2 (DML2) and DML3 through a base excision repair pathway (11, 12). However, epigenetic profiles are dynamic in response to biotic and abiotic stress, and during reproduction and development. Similar to mammals, complete epigenetic reprogramming is required during flowering plant gamete formation, characterized by extensive DNA demethylation, which in Arabidopsis is performed by DME (13, 14).

DME encodes a bifunctional 5mC DNA glycosylase/lyase that is essential for reproduction (11). Paralogs ROS1, DML2, and DML3 function primarily in the sporophyte to counteract the spread of DNA methylation mediated by RNA-dependent DNA methylation (RdDM) (15, 16). The A, Glycosylase, and B regions of the C-terminal half of DME are conserved among the DME/ROS1 DNA glycosylase clade, and are absolutely required for DME 5mC excision activity in vitro, comprising the minimal catalytic core for direct excision of 5mC from DNA (11, 17, 18). DME acts specifically in the central cell of the female gametophyte and the vegetative nucleus of pollen (17, 19, 20). The pollen grain contains the vegetative nucleus and two sperm cells, and the vegetative nucleus contributes to the germination and growth of the pollen tube, which delivers the to sperm cells to the female gametophyte. Following double fertilization, the egg and central cell respectively develop into the embryo and nutritive endosperm, which accumulates starch, lipids, and storage proteins to nourish the developing embryo (21, 22). The endosperm is also the site of plant genomic imprinting, i.e. parent-of-origin specific expression, resulting from allelic inheritance of differential epigenetic states. DNMT1 homolog MET1-mediated DNA methylation and DME-mediated demethylation are primary players in the epigenetic regulation of endosperm imprinting (11, 23–26). For example, DME-mediated DNA demethylation is required for the activation of *MEA, FIS2*, and *FWA* expression in the central cell, which persists in the endosperm, while MET1 is responsible for the silencing of *FIS2* and *FWA* paternal alleles (23, 25). Imprinting is essential for reproduction in Arabidopsis, and seeds that inherit a maternal *dme* allele abort due to failure to activate *MEA* and *FIS2*, essential components of the endosperm Polycomb Repressive Complex 2 (PRC2) required for seed viability (17, 23).

Although DME preferentially targets small, AT-rich, and nucleosome-poor euchromatic transposons, it also demethylates intergenic and heterochromatin targets (13). How DME is recruited to distinct target sites with various chromatin structures is largely unknown, although the <u>FA</u>cilitates <u>C</u>hromatin <u>T</u>ransactions (FACT) histone chaperone is required at heterochromatic targets and some imprinted loci (27, 28). Other than the glycosylase domain, the catalytic core region of the DME/ROS1-like DNA glycosylases contains multiple conserved globular domains of unknown function.

Here, we show that expressing a nuclear-localized DME C-terminal catalytic region controlled by a native DME promoter complements *dme* seed abortion and pollen germination defects, and partially rescues the DNA hypermethylation phenotype in endosperm. DNA methylation analysis revealed that the majority of canonical DME target sites are demethylated by the catalytic core, indicating that this region is sufficient to direct DME targeting and localization. However, without the N-terminal region, the degree of demethylation is reduced across all target sites and demethylation of heterochromatin targets is particularly impeded. In addition, we observed prevalent ectopic demethylation specifically at genic sequences by the catalytic core. Thus, the N-terminal region of DME is needed for the full breadth and depth of demethylation and to prevent gene body demethylation. We provide evidence that the N-terminal conserved domains are specific to the angiosperm lineage, acquired late during land plant evolution, potentially to ensure robust demethylation in nucleosome-rich heterochromatin targets.

## Results

### Nuclear-Localized Catalytic Core of DME Rescues *dme* Associated Developmental Defects

*DME* encodes at least two alternatively spliced variants, capable of producing two hypothetical polypeptides of 1,729 (DME.1) and 1,987 (DME.2) amino-acids in length (17, 29). The amino acid positions denoted in this study correspond to DME.2, the predominant isoform expressed in floral tissues (19). First, we detailed the domain structure and predicted characteristics of the catalytic and non-catalytic regions of the DME protein. The N-terminal half of DME consists of a large portion of unstructured, low complexity sequences (aa 364-947, N-terminal variable region, Fig. 1A), a stretch of basic amino acid-rich repeats (aa 291-363, Basic stretch) previously shown to direct nuclear localization (17), and an 120 amino-acid N-terminal domain (aa 1-120, DemeN) of unknown function present only among the angiosperm DME/ROS1-like proteins. Within the DemeN domain is a sWIPxTPxKs motif that is highly conserved (Fig. S1) but is absent in the shorter DME.1 isoform. This motif has a hydrophobic core and may mediate protein-protein interactions and/or contribute to DME regulation by post-translational modification i.e. phosphorylation or methylation at the lysine residue. The basic stretch region is highly conserved among angiosperm DME-like proteins and is reminiscent of the AT-hook motifs that can bind DNA in a non-sequence specific manner (30), suggesting the basic stretch might also bind DNA, in addition to directing DME to the nucleus.

**Fig. 1.**
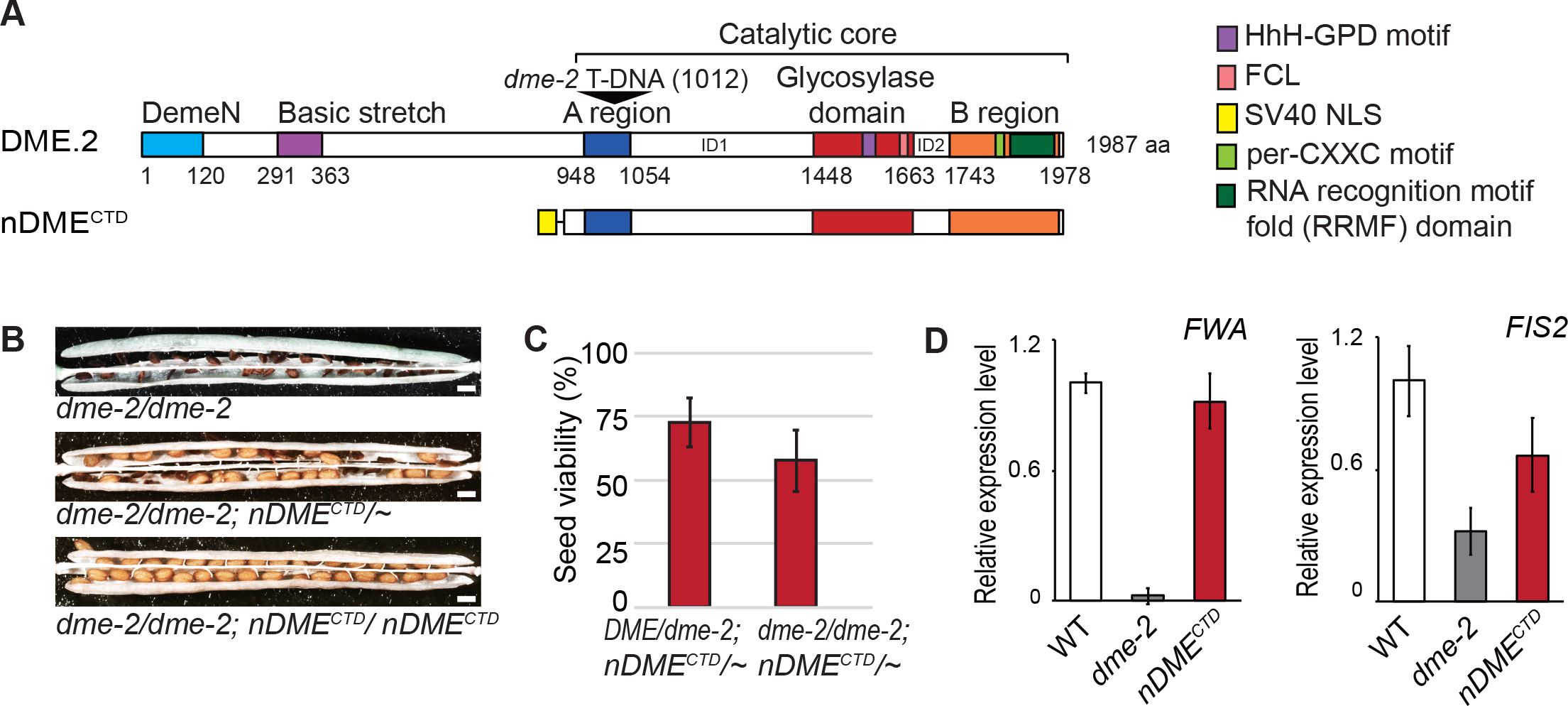
Complementation results. (A) Domain architecture and the positions of conserved domains along DME protein. nDME^CTD^ is the construct used for complementation and methylome analyses. (B) In *dme-2/dme-2* siliques > 99% of seeds are aborted. A single copy of the *nDME^CTD^* transgene reduces seed abortion to 50%; and in the *dme-2/dme-2*; *nDME*^*CTD*^*/nDME*^*CTD*^ siliques, all seeds are rescued and develop normally. Scale bar = 0.5 mm. (C) The percentages of viable seeds in *DME/dme-2* or in *dme-2/dme-2* plants that were complemented by *nDME*^*CTD*^ transgene. Error bars represent standard deviations (D) The *nDME*^*CTD*^ transgene restores DME target genes *FWA* and *FIS2* expression. WT: Col-0; nDME^CTD^: *dme-2/dme-2*; *nDME*^*CTD*^*/nDME*^*CTD*^; *dme-2*: *dme-2* homozygotes. Total RNA was isolated from stage F1 to F12 floral buds.

It was previously determined that the C-terminal half of DME (aa 936-1987, hereafter the DME^CTD^) constitutes the catalytic core for 5mC excision in vitro (11, 31). This core contains the catalytic glycosylase domain of the HhH (<u>h</u>elix-<u>h</u>airpin-<u>h</u>elix) motif followed by the [4<u>F</u>e-4S] <u>c</u>luster <u>l</u>oop (FCL) motif (32). The conserved B region contains a domain predicted to adopt an <u>R</u>NA <u>R</u>ecognition <u>M</u>otif-like fold (RRMF) and a divergent, circularly permuted version of a singlecopy methylated CpG-discriminating CXXC domain (32). The presence of a permuted CXXC and RRMF raises the possibility that the enzymatic core might contain intrinsic targeting information (33, 34). To test this possibility, we investigated whether DME^CTD^ can rescue *dme* seed abortion phenotype when expressed by an endogenous *DME* promoter. A classical SV40 NLS (PKKKPKV) was included to ensure robust nuclear localization (Fig. 1A). The resulting transgene (*nDME^CTD^)* was transformed directly into *DME/dme-2* heterozygous Col-g/ plants via the floral dipping procedure (35). Self-pollinated non-transformed *DME/dme-2* plants produce 50% viable (inherited *DME* maternal allele) and 50% aborted (inherited *dme-2* maternal allele) F1 seeds. In contrast, transgenic lines that were *DME/dme-2* and hemizygous for a single locus *nDME^CTD^* transgene, produced a 3:1 ratio of viable to aborted F1 seeds when self-pollinated (1292:439, 3:1, χ^2^ = 0.12, P > 0.7, Table S1), indicating that *nDME^CTD^* fully complemented the dme-mediated seed abortion phenotype. When the *nDME^CTD^* transgene was transformed into *dme-2/dme-2* homozygous plants (Materials and Methods), the resulting T1 transgenic lines (*dme-2/dme-2;nDME^CTD^/~*) displayed 50% to 75% viable seed rates when self-pollinated (Table S1), compared to selfed non-transformed *dme-2/dme-2* plants which bear < 0.05% viable seeds, again indicating full seed abortion complementation by *nDME^CTD^* (Fig. 1B, C, Table S1). These results show that nDME^CTD^ can rescue the *dme-2* seed abortion phenotype. Since *dme* mutant seed abortion is partially due to defects in activating imprinted *Polycomb Repressor Complex 2* (*PRC2*) genes (36), we used qRT-PCR to measure expression of PRC2 subunit *FIS2*, and *FWA*, whose maternal expression is enabled by DME-mediated DNA demethylation, and found that nDME^CTD^ restored the expression of these genes (Fig. F1D).

DME is also expressed in the vegetative cell of pollen, and mutations in *DME* reduce pollen germination, resulting in lower transmission of the paternal mutant *dme* allele in certain ecotypes (20). Under our growth conditions, when *DME/dme-2* heterozygous Col-*gl* plants were self-pollinated, about 20-30% (compared to the expected 50% if *dme-2* allele is normally transmitted) of the F1 progeny were *dme-2* heterozygotes (Table S2). To test whether nDME^CTD^ can rescue the *dme* pollen phenotype, we pollinated wild-type Col-0 with pollen derived from transgenic lines that are homozygous for the *dme-2* allele and carry a single locus of the *nDME^CTD^* transgene (*dme-2/dme-2; nDME^CTD^/~* lines with ~50% seed abortion rates, Table S1). If nDME^CTD^ does not complement *dme-2* pollen germination defects, we expect roughly half of the F1 progeny will carry the nDME^CTD^ transgene (hygromycin resistant) because mutant pollen with or without the transgene would germinate with equal frequency. Instead, we observed 65% - 90% of the F1 progeny are hygromycin resistant (resistant: sensitive = 190:52, 1:1, χ^2^ = 79.69, P = 7.3E^−19^; Table S3), indicating that *nDME^CTD^* rescues *dme-2* pollen defects. These results show that nDME^CTD^ can rescue the *dme* developmental phenotype, which indicates that DME targeting and recruitment information is contained within the catalytic core region.

### Canonical DME target loci are demethylated by nDME^CTD^

The molecular cause of DME mutant phenotypes are a loss of DME-mediated genome-wide DNA demethylation. To test the extent of nDME^CTD^ complementation, we compared the methylome of the nDME^CTD^-complemented endosperm to those of wild-type and *dme-2* endosperm (13). nDME^CTD^-complemented endosperm methylomes from three independent lines (*dme-2/dme-2*, *nDME*^*CTD*^*/nDME*^*CTD*^, F2 and F3 generation of *dme-2* homozygotes) were generated and the reads combined for downstream analyses (Pearson correlation coefficients between independent lines showed they were highly concordant >0.93 (Table S4)). We compared the differentially methylated regions (DMRs) between *dme-2* and wild-type endosperm (DME targets), and the DMRs between *dme-2* and nDME^CTD^-complemented endosperm (nDME^CTD^ targets) (Materials and Methods) (13). Looking genome-wide, but excluding all DME and nDME^CTD^ targets, the Pearson correlation coefficients between our combined independent lines and the previously published wild-type and *dme-2* endosperm datasets ranged between 0.92 and 0.94 (Table S5), indicating a high level of concordance between the methylomes used in this study.

We found several DME-regulated imprinting control regions of maternally and paternally expressed genes (*MEGs* and *PEGs*, respectively) to be hypomethylated in the *nDME^CTD^*-complemented endosperm compared to *dme-2* endosperm (Fig. S2), suggesting that nDME^CTD^ is active at these loci. CG methylation does not return to wild-type levels, however, indicating that the genome is demethylated to a lesser degree by nDME^CTD^ than by wild-type DME (Fig. 2A and Fig. S3C). The DME and nDME^CTD^ DMRs largely overlap (Fig. 2B), and for the DMRs that appear unique to DME, the same regions are also demethylated by nDME^CTD^ (Fig 2C, black solid-line trace), but to a reduced degree (the solid black peak is on the left of the dotted peak) that falls below our DMR cutoff (fractional CG methylation difference ≥ 0.3, Fisher’s exact test *p*-value < 10^−10^). The shared DMRs are also slightly less demethylated in nDME^CTD^-complemented endosperm compared to wild-type endosperm (Fig. 2D, red trace is to the left of the black trace). Together, these data show that nDME^CTD^ rescues the *dme*-associated endosperm hypermethylation phenotype, but only partially. This reduced demethylation was not a result of low transgene expression as far as we could ascertain, since qRT-PCR analyses of endosperm tissue showed that the *nDME^CTD^* is abundantly expressed (Fig. S2D).

**Fig. 2.**
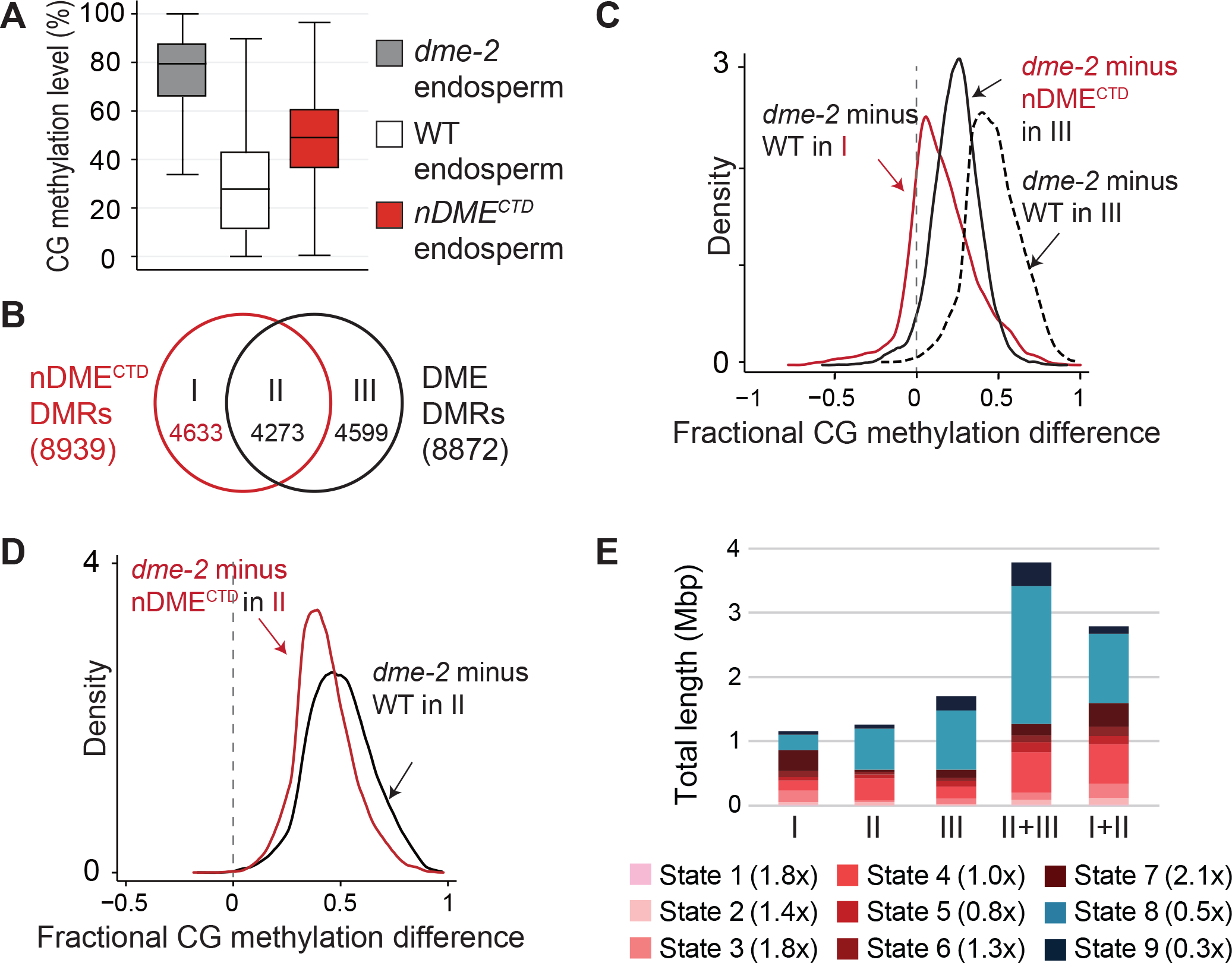
Endosperm methylome analysis. (A) Boxplot of CG methylation levels among canonical DME target sites in *dme-2* mutant (grey), wild-type (white), or in *nDME^CTD^*- complemented (red) endosperm. (B) Venn diagram depicting unique and shared regions and loci numbers between DME and nDME^CTD^ DMRs. (C) Kernel density plots of CG methylation differences between *dme-2* and WT endosperm (black dotted trace), or between *dme-2* and nDMECTD-complemented endosperm (black trace), for DME unique DMRs, and CG methylation difference between *dme-2* and WT endosperm, for nDME^CTD^-unique DMRs (orange trace). (D) Kernel density plots of CG methylation differences between *dme-2* and *nDME*^*CTD*^-complemented endosperm (black trace) or CG methylation differences between *dme-2* and WT endosperm (orange trace), within the DME and nDME^CTD^ shared DMRs. (E) Chromatin state distribution, and total length covered, within nDME^CTD^-unique (I), nDME^CTD^-DME shared (II), DME-unique (III) DMRs, DME-all (II+III), and nDME^CTD^-all (I+II) DMRs. State 1 to 7 correspond to euchromatin and states 8 to 9 correspond to AT- and GC-rich heterochromatin, respectively. The numbers in the brackets show fold changes (total length covered) in nDME^CTD^ relative to DME DMRs.

To investigate whether chromatin features influence nDME^CTD^ recruitment and demethylation, we assessed the specific histone marks and genomic characteristics (37) in target sites that are nDME^CTD^-unique, DME-unique, or shared between the two. Compared to nDME^CTD^-DME shared DMRs, DME-unique target sites are highly enriched for heterochromatin states 8 and 9 (Fig. 2E). nDME^CTD^ DMRs (unique and shared) are enriched for chromatin states 1, 6, and 7 that correspond to open chromatin states, but show a significant reduction in heterochromatic chromatin states 8 and 9 when compared to DME DMRs (Fig. 2E)(37). Thus, nDME^CTD^ demethylates poorly at heterochromatic loci and preferentially targets euchromatin sites.

### The DME N-Terminal Region is Required for Effective Demethylation at Long Heterochromatic Target Sites

Longer DME DMRs almost exclusively reside in heterochromatin (86.3% of 1-1.5 kb and 95.5% of ≥ 1.5 kb, Fig. 3A). We postulate that this is due to the dense methylation associated with heterochromatin, which may result in longer stretches of DNA demethylation during DME occupancy at these sites. Interestingly, the number of long DMRs is dramatically reduced in nDME^CTD^-complemented endosperm (Fig. 3A, B). This reduction in the number of longer DMRs was not due to a lack of nDME^CTD^ targeting to these sites, since the partial demethylation characteristic of nDME^CTD^ activity occurred in all targets regardless of their length (Fig. S3E). However, when we analyzed the length of the nDME^CTD^ demethylated regions, we found that it produced much shorter DMRs (Fig. S5). For example, there are 250 DME DMRs that are longer than 1.5 kb (median = 1.9kb). Among them, 165 are also DMRs of nDME^CTD^, but are much shorter in length (median = 400bp) (Fig. 3C-D). Thus, removal of the DME N-terminal region significantly reduced the extent of demethylation in these long targets.

**Fig. 3.**
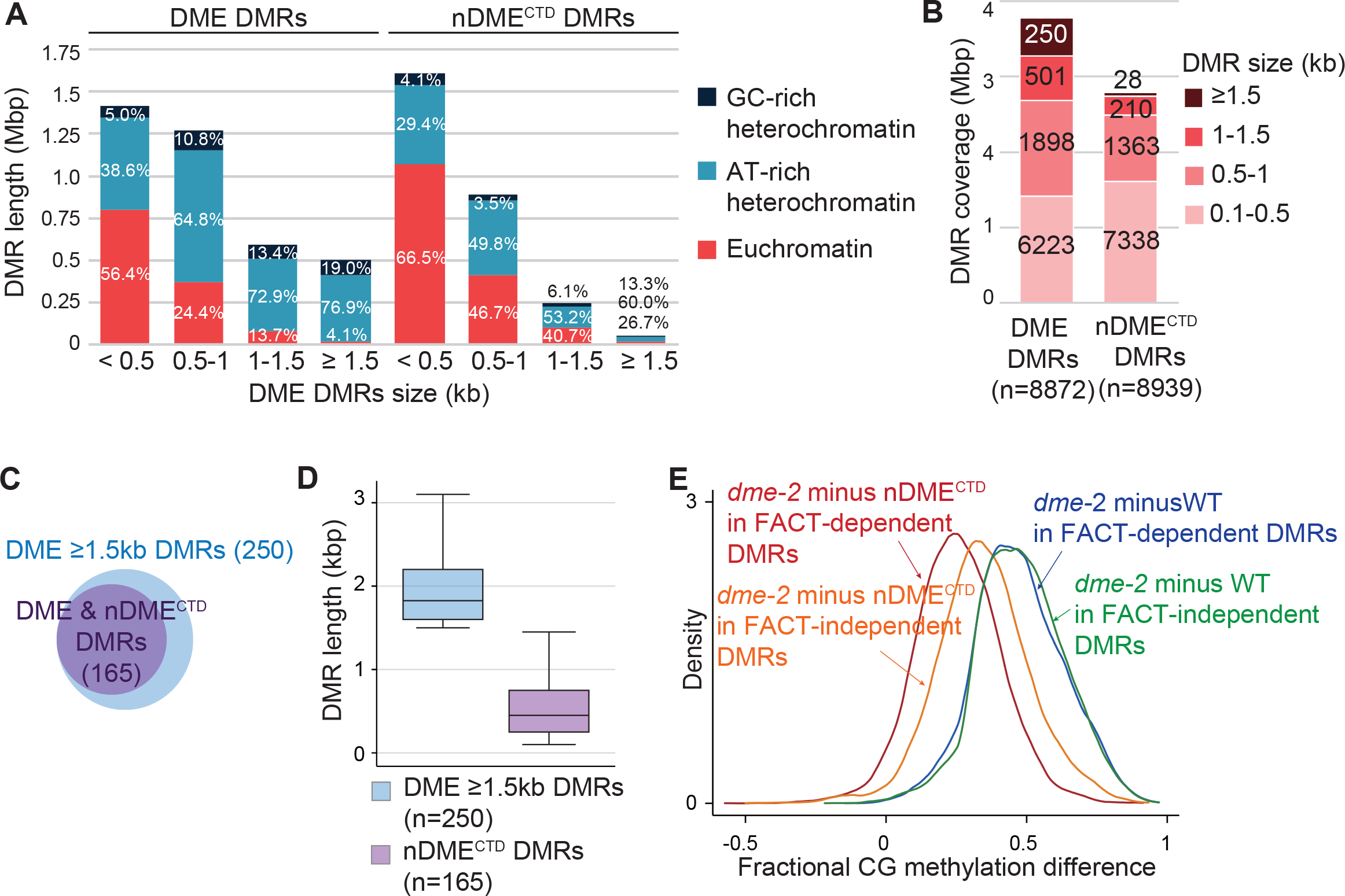
DME and nDME^CTD^ DMRs. (A) Distribution of euchromatin and heterochromatin within each DMR length group. (B) DME and nDME^CTD^ DMRs grouped by their sizes, with the total length they cover shown. (C) The majority of the 250 longer DME DMRs overlap with the DMRs of nDME^CTD^, but the nDME^CTD^ DMRs are much shorter in size (D). (E) FACT dependency of the DME targets demethylated by DME (FACT-dependent, dark green trace; FACT-independent, light green trace) or by nDME^CTD^ (FACT-dependent, magenta trace; FACT-independent, orange trace).

The histone chaperone FACT complex is required for demethylation of about half of DME targets in Arabidopsis, particularly those in longer TEs in heterochromatin regions (27). DME co-localizes with SPT16 (the larger of the two FACT subunits) in an in vivo BiFC assay, suggesting that DME might recruit the FACT complex to these heterochromatic target loci (27). Of the 250 long DME DMRs, 87% of them require FACT activity, raising the possibility that nDME^CTD^ might be defective in recruiting FACT. To test this hypothesis, we examined how nDME^CTD^ demethylates FACT-dependent vs independent loci. In wild-type endosperm, both target groups are demethylated to a similar degree (Fig. 3E, blue and green traces have similar shape and peak). In the nDME^CTD^-complemented endosperm, FACT-independent loci are only slightly less demethylated compared to wild-type endosperm (Fig. 3E, orange trace moderately shifted to the left of blue and green traces). In contrast, demethylation at the FACT-dependent loci is more severely impeded (Fig. 3E, red trace). These results support a model whereby DME recruits FACT via the N-terminal region to make heterochromatic regions accessible.

### Prevalent Ectopic Gene Body Demethylation by nDME^CTD^

Our analyses revealed a set of new DMRs that were unique to nDME^CTD^, which we term as ‘ectopic’ targets (Fig 2B). The CG methylation difference between *dme-2* and wild-type endosperm for these DMRs is minimal but not absent (Fig. 2C, although the red trace peak is close to zero, there is a positive right-hand shoulder). As shown in Fig. 4A, we plotted the methylation status of nDME^CTD^-unique loci in wild-type endosperm to assess how they are demethylated by DME. This resulted in positive peaks for shorter (red trace) or longer (blue trace) transposable elements (TEs), and intergenic sequences (green trace), showing that these nDME^CTD^-unique loci are also demethylated by DME but not enough to reach the DMR cutoff (fractional CG methylation difference ≥ 0.3, Fisher’s exact test p-value < 10^−10^). In contrast, most nDME^CTD^-unique loci within coding sequences are not demethylated by DME (Fig. 4A, orange trace). This indicates that genic sequences are the primary ectopic targets of nDME^CTD^. This is also reflected by the increased nDME^CTD^ DMR frequency (Fig. 4B, red and orange traces), the decreased average CG methylation (Fig. S4) within coding genes. These results show that nDME^CTD^ preferentially targets methylated coding genes.

**Fig. 4.**
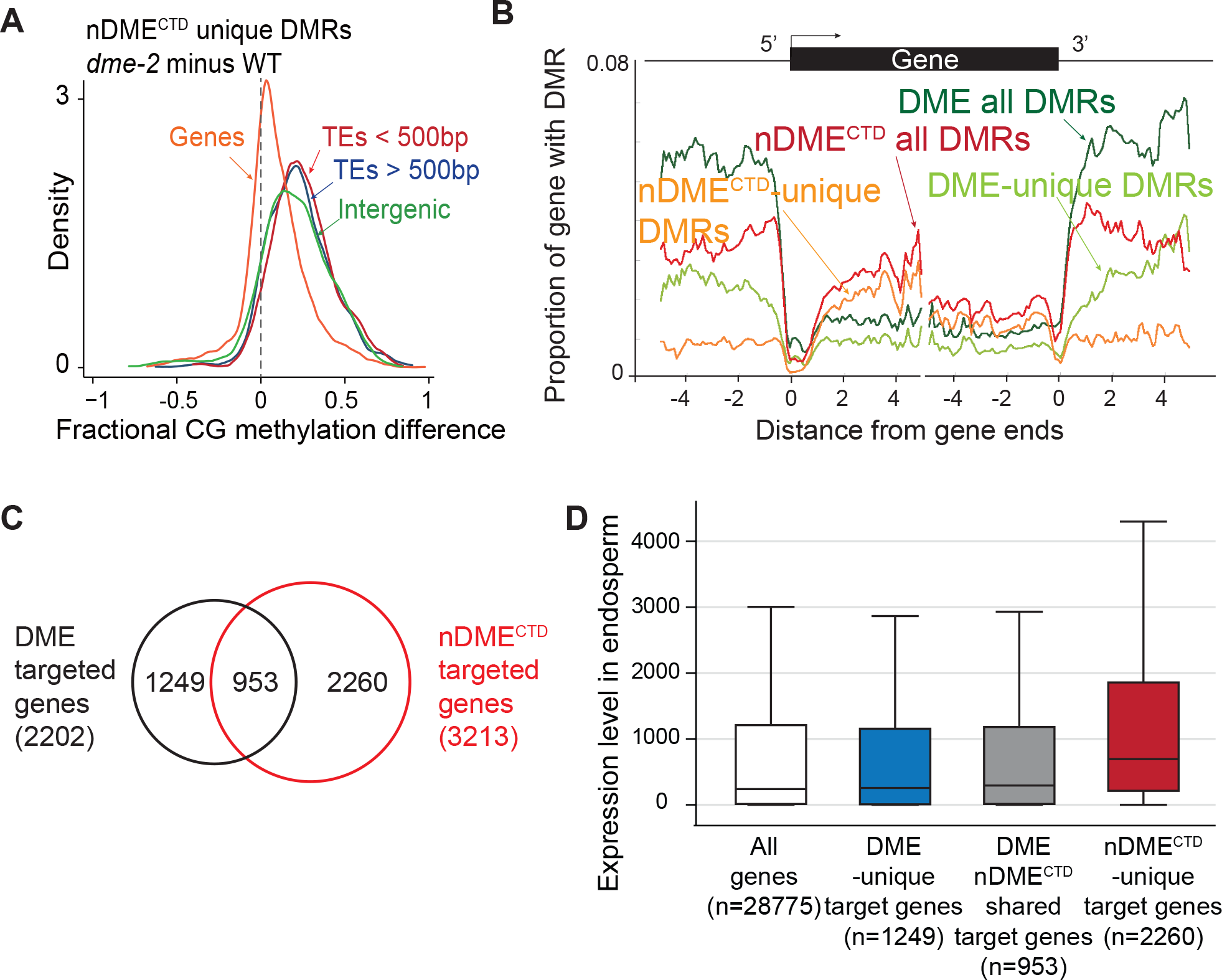
nDME^CTD^ induces ectopic genic demethylation. (A) Kernel density plot of CG methylation differences between *dme-2* and wild-type endosperm, for nDME^CTD^-unique loci that reside in short (< 500bp, red trace) or longer (>= 500bp, blue trace) TEs, intergenic regions (green trace), or genic sequences (orange trace). (B) Distribution frequency of DMRs with respect to coding genes. Genes were aligned at the 5’ end or the 3’ and the proportion of DMRs in each 100-bp interval is plotted. DMR distribution is shown with respect to all DME DMRs (dark green trace), DME-unique DMRs (light green trace), all nDME^CTD^ DMRs (red trace), and nDME^CTD^-unique DMRs (orange trace). (C) Venn diagram showing the numbers of coding genes associated with DME and nDME^CTD^ DMRs. (D) Transcriptional scores (RPKM) in wild-type endosperm using expression data from (13, 14) for each group of genes indicated.

Gene body CG methylation, or gbM, is an evolutionarily conserved feature in mammals and angiosperms, but the origin of gbM remains unknown and its function is highly debated (3, 4). Around 15% (~5,000) of Arabidopsis genes contain gbM (38). In our endosperm methylomes, we found 2,202 and 3,213 genes are associated with DME or nDME^CTD^ DMRs, respectively, which were largely mutually exclusive and thus constituted almost all of the genes with gbM (Fig. 4C). Among them, 2,260 genes were ectopically targeted by nDME^CTD^ (Fig. 4C, Dataset S1). These nDME^CTD^-unique genes have higher CG methylation and higher expression levels compared to DME-targeted coding genes (Fig. 4D). They include genes across most actively used cellular processes (Table S6), consistent with the current theory that moderate gene body methylation positively correlates with constitutively transcribed genes (3). These observations suggest the presence of an active mechanism that prevents DME from targeting genes undergoing active transcription in wild-type plants, but which is impaired by nDME^CTD^.

### The Evolutionary History of the DME/ROS1 Glycosylase Family

Our data show that the C-terminal catalytic region of DME is able to efficiently demethylate DNA in vivo, and also harbors targeting ability. We also demonstrate that the N-terminal domain is important for fine-tuning DME targeting to heterochromatin and restricting it from gene bodies. To provide clues as to the evolutionary origin of these bipartite domains, we carried out comparative analyses across plant lineages. Using various DME homologs as query sequences, we revealed a diversity of N-terminal domains associated with the DME catalytic core across various clades (Fig. 5). These indicated that a shorter protein, comprising only the C-terminal of Arabidopsis DME, may represent the ancestral form of the 5mC glycosylase that is found in all plant lineages. Variation of the N-terminal domains included land plants and charophytes (streptophyta), which possess a divergent circularly permuted CXXC domain between the FCL and RRMF domains. In contrast, one or more copies of the classical CXXC can be found in chlorophyte and stramenopile algae at distinct positions. Chlorophyte and stramenopile homologs also possess additional chromatin readers such as Tudor and PHD, DNA-binding domains such as the AT-hook domain, and the chaperone Hsp70-interacting DnaJ domains (32, 39). These accessory domains suggest a mode of regulating DNA glycosylase activity according to methylation status (via CXXC) or chromatin states (via PHD, Tudor). The DemeN and basic-stretch of the DME N-terminal region are restricted to the angiosperm lineage and appear to be a late acquisition during land plant evolution (Fig. 5). The acquisition of this region thus coincides with the origins of double fertilization in plants and emergence of plant gene imprinting.

**Fig. 5.**
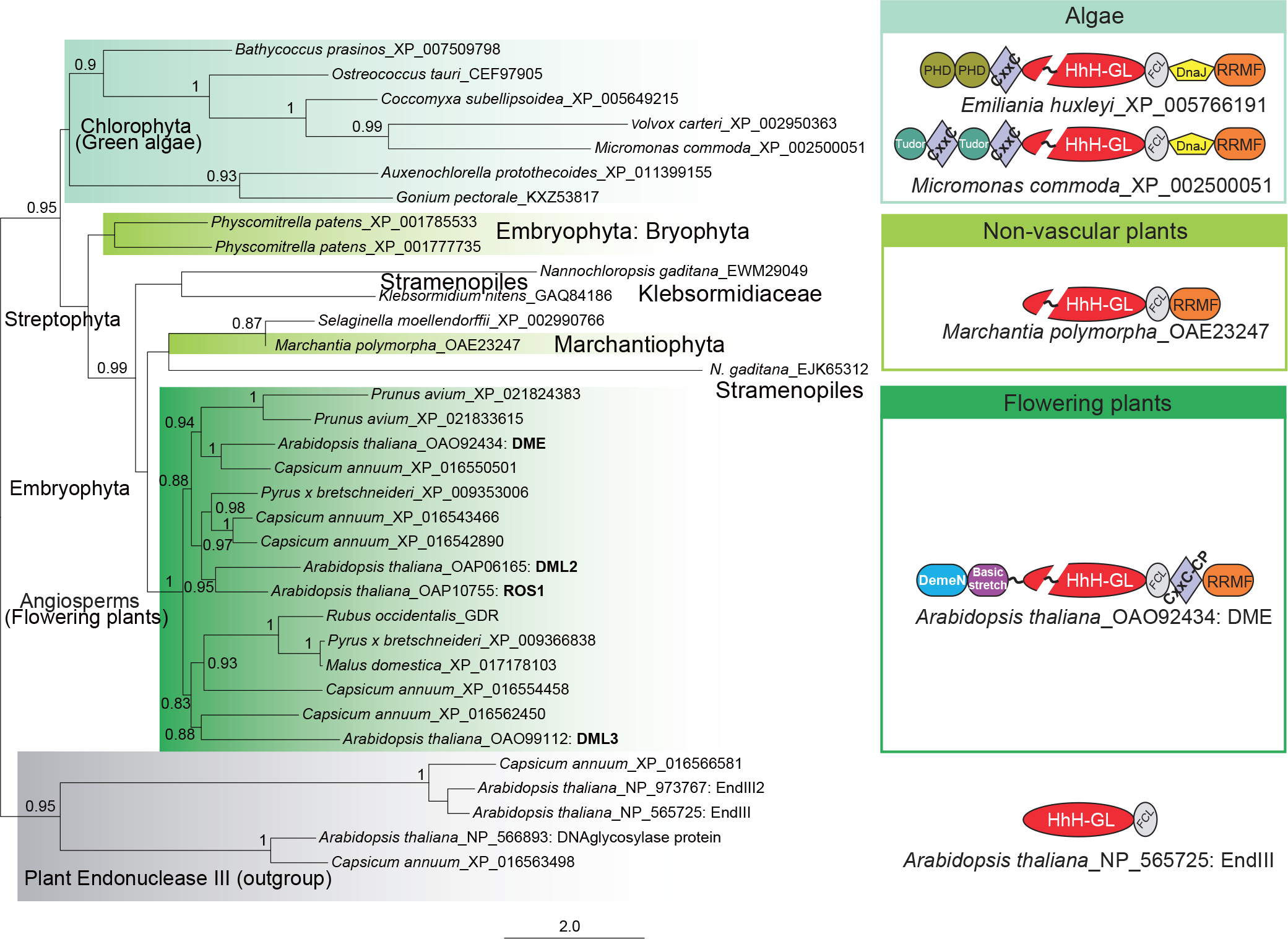
Evolution of plant DME-like proteins. A phylogenetic tree was reconstructed using the PhyML program. Only node supporting values >0.80 from ML bootstrap analyses are shown. The representative domain architectures of DME homologs in major plant clades are shown along the tree, demonstrating domain fusions during evolution. Domain abbreviations: DemeN, N-terminal domain of DME-like proteins in angiosperms; DnaJ, DnaJ molecular chaperone homology domain (Pfam: PF00226); FCL, [Fe4S4] cluster loop motif (also called Iron-sulfur binding domain of endonuclease III; Pfam: PF10576); HhH-GL, HhH-GPD superfamily base excision DNA repair protein (Pfam: PF00730); PHD, PHD finger (Pfam: PF00628); RRM, RNA recognition motif (Pfam: PF00076); Tudor, Tudor domain (Pfam: PF00567).

## Discussion

DME is an essential epigenetic modifier that regulates gene imprinting and influences trans-generational epigenetic inheritance in Arabidopsis (40). DME demethylates the central and vegetative gamete companion cell genomes at thousands of loci, but the mechanism of DME targeting remains elusive. This is due to its restricted expression to these ephemeral nuclei, embedded within Arabidopsis gametophytes and largely prohibiting biochemical interrogation, at least with current tools. In contrast, genetic analysis, coupled with endosperm transcriptome and methylome profiling, has been instrumental in revealing DME’s molecular function (11, 13, 14, 27, 41). Here, we used genetic complementation and endosperm DNA methylation profiling assays to show that the catalytic core of the DME protein, containing the DNA glycosylase, FCL, permuted CXXC and divergent RRMF domains, is sufficient to rescue the *dme* seed abortion phenotype and complement the pollen germination defects caused by the *dme* mutation (Fig. 1). We also present evidence that nDME^CTD^ can be recruited to, and demethylate, most canonical DME target sites, and reason that the DME C-terminal catalytic region must contain sufficient information for targeting and recruitment. We go on to suggest the DME protein has a bipartite structure (Fig. S7) and demonstrate a requirement for the N-terminal domain in targeting DME to heterochromatin, possibly via FACT.

Although our nDME^CTD^ complemented *dme*-associated developmental defects (Fig 1B, Table S1, S3), it did not fully rescue the *dme* endosperm DNA hypermethylation phenotype. Instead, we observed a reduced degree of demethylation by nDME^CTD^ in all of the endogenous DME target loci, regardless of their length (Fig. 2A, S2E). From this, we deduce that the DME N-terminal region is likely required for full and robust demethylation activity. To achieve this, the N-terminal region may assist the glycosylase by tightly binding to DNA templates (via the AT-hook) to promote complete and thorough demethylation. Previous studies of ROS1 support such a model: the basic stretch/AT-hook region of ROS1 was shown to bind strongly to DNA templates in vitro in a non-sequence-specific manner, and removal of the ROS1 basic-rich region impaired the sliding capacity of ROS1 on the DNA template (42), significantly reducing ROS1 5mC excision activity (30).

DME preferentially targets smaller euchromatic transposons that flank coding genes, it also targets gene-poor heterochromatin regions for demethylation (13). Since heterochromatin regions are compacted, demethylation in these regions requires substantial chromatin remodeling, including the temporary eviction of nucleosomes in order for DME to gain access to DNA. This is reminiscent of the requirement for the ATP-dependent chromatin remodeler DDM1 at linker histone H1-associated heterochromatin for methyltransferases access (43). The nucleosome remodeler FACT was previously shown to be required for DME-mediated demethylation, primarily at heterochromatin targets, including those associated with H1 (27), and we additionally noted that these sites have increased nucleosome occupancy (Fig. S6). It is tempting to speculate that the DME N-terminal region is required to recruit factors such as FACT, and indeed we found the overwhelming majority of the 250 longer DME DMRs (87%) that were not demethylated in the absence of the N-terminal region, also require FACT activity for demethylation (Dataset S2). SPT16 was shown to co-localize with DME in vivo (27), suggesting that a direct or indirect interaction between DME and FACT does occur.

nDME^CTD^ also displayed a reduced capacity in demethylating FACT-independent loci. Thus, it is possible that the N-terminal region is needed to recruit other chromatin remodelers at FACT-independent targets, i.e. if nucleosomes are a natural barrier for DME demethylation in euchromatin as well as heterochromatin (Fig. S6). We envision a working model (Fig. S7) where the catalytic region is sufficient for directing DME to target sites, whilst the N-terminal region is required to interact with the local chromatin environment, stabilizing binding to the chromosomal template, and/or assisting demethylation of flanking sequences by remodeling nucleosomes.

Our observation that nDME^CTD^ demethylated a large number of ectopic sites in genic sequences was unexpected. Gene body methylation is evolutionarily conserved and around 20% of all Arabidopsis genes contain gbM (38, 44–47). Genes that are CG-methylated are preferentially constitutively expressed housekeeping genes (48), raising the possibility that these genic methylated genes reside in open chromatin regions, easily accessible by chromatin or DNA modifying enzymes, so that nDME^CTD^ was able to act in those regions, whereas the more tightly-regulated endogenous DME is not. It is also possible that DME is actively repelled by certain open chromatin histone marks such as H3K4me1, and the region responsible for repulsion is missing in nDME^CTD^. This scenario would be analogous to the mammalian de novo DNA methyltransferase DNMT3, where its binding to an allosteric activator, unmethylated histone H3, is strongly inhibited by H3K4 methylation (49, 50).

Tracing the evolutionary history of the DME-like genes (Fig. 5), we found that a bacterial version of the HhH-FCL pair underwent a horizontal gene transfer to the ancestor of plants, followed by a gene duplication. One copy was fused to an RRMF domain and further acquired an insert in the glycosylase domain, giving the ancestral form of DME in plants. This was likely then transferred to the stramenopiles from a secondary chlorophyte endosymbiont of this lineage. Finally, at the base of the streptophyte radiation, DME acquired a permuted CXXC, and later the DemeN domain and associated charged repeats were acquired in angiosperms, possibly to facilitate and ensure robust and thorough DNA demethylation. Thus, the adoption of a DME-based demethylation system for DNA base modification appears to have occurred early in the plant lineage. The presence of several accessory domains in addition to the conserved core suggests variation in the chromatin environments in specific lineages. For example, the presence of the DemeN and basic stretch/AT-hook motifs in angiosperms and the permuted CXXC domain in the streptophyta lineage likely reflect adjustment to the unique methylation and chromatin environment of the larger Streptophyta and land plant genomes.

In summary, our data show that the catalytic core of DME was present in ancient plant ancestors, and is alone capable of targeting and demethylation. We provide evidence that the N-terminal domain, confined to flowering plant DME proteins, evolved to facilitate access to diverse chromatin states, in turn mediating gene imprinting and the transgenerational silencing of transposons (13).

## Materials and Methods

Complete details concerning plant materials, experimental methods, and data analyses are provided in the Supplemental Information.

## Author contributions

C.Z., Y.-H.H., R.L.F. and T.-F.H. designed research; C.Z., Y.-H.H., X.-Q.Z., and F.L. performed research; C.Z., Y.-H.H., X.-Q.Z., D.Z., J.M.F., L.M.I., L.A., and T.-F.H. analyzed data; J.M.F, R.L.F. and T.-F.H. wrote the paper, and W.X., L.M.I., L.A., J.H.H. provided insightful comments on the research and the manuscript.

## Acknowledgements

We thank J.-Y. Lin (UCLA), P.-H. Hsieh (JGI), D.B. Lyons (UC Berkeley), and Y.H. Choi (Seoul National University) for suggestions and critical reading of the manuscript. This work is supported by the Hatch Project (No. 02413) of National Institute of Food and Agriculture, USDA to T.-F.H., the National Science Foundation Grant MCB-1715115 to T.-F.H. and W.Y.X. LMI and LA are supported by the IRP funds of the National Institutes of Health, USA.

## Data availability

All data generated or analyzed in this study are included in this article and Supplementary Information files. BS-seq sequencing data are available in NCBI (GEO XXXXXX).

