## Supplemental Information for "The Catalytic Core of DEMETER Guides Active DNA Demethylation in Arabidopsis"

**This PDF file includes:**

Supplementary text
Figs. S1 to S7
Table S1 to S6
References for SI reference citation

**Other supplementary materials for this manuscript including the followings:**

Dataset S1
Dataset S2

**Zhang el al., Supplemental Information**

**Materials and Methods**

**Construction of transgenes used in this study.**

A binary plasmid vector, pFGAMh, was modified to facilitate the generation of plasmid constructs using the Gibson assembly method. In brief, the pFGAMh vector backbone was derived from pFGC5941 (GenBank Accession: AY310901). In pFGAMh, the T-DNA region of pFGC5941 was replaced by a hygromycin resistance cassette as a plant selectable marker and a Gateway attR cassette (rfa, Invitrogen, Carlsbad CA, USA). The hygromycin resistant gene (HPTII) was driven by a mannopine synthase (MAS) promoter and terminated by a MAS terminator. The Gateway attR cassette was flanked with XhoI and XbaI unique restriction sites and followed by an octopine synthase (OCS) terminator.

The pDME: nDME$^{CTD}$ plasmid comprises a linearized pFGAMh vector and three DNA fragments: a DME regulatory sequence (DMEpro), a bridge sequence, and the DME$^{CTD}$ coding sequence (linker-DME$^{CTD}$). Vector plasmid pFGAMh was linearized by digesting with restriction enzymes XhoI and XbaI and removing Gateway attR cassette. The first fragment contains 2895 bp upstream of the DME.2 translation start codon ATG (DME regulatory sequence; DMEpro), which was PCR-amplified from Col-0 gDNA with primer pair VeDME/P3R. The second fragment was generated by annealing DNA oligos S40F and S40R that contains the Kozak consensus sequence (AACAATGGTG), SV40 NLS (CCAAAGAAGAAGAGAAAGGTC). The third fragment was PCR-amplified from Col-0 cDNA with primer pair lnAGBF/CTDVeR. This fragment contains a 6-Alanine linker sequence and 3159 bp of C-terminal DME coding sequence (linker-DME$^{CTD}$, total 3174 bp). The linearized pFGAMh vector and the three DNA fragments were assembled using Gibson assembly (NEB) to generate pDME:nDME$^{CTD}$. Primer sequences are listed in Table S7.

**Plant materials and seed phenotype analysis.**

Heterozygous *DME*/*dme-2* lines in the Col-*gl* background were subjected to Agrobacterium-mediated transformation. Seeds were sterilized by 30% bleach solution and plated on a 0.5x MS nutrient medium with 1% sucrose, 0.8% agar and 50 $\mu$g/ml kanamycin, and stratified at 4°C for 2 days. Germinated seedlings were transferred to soil and grown in the growth room under 16 hours of light and 8 hours of dark cycles at 23°C. Siliques from $T_1$ transgenic plants were dissected 14-16 days after self-pollination using a stereoscopic microscope (SteREO Discovery.V12, Carl Zeiss, Wetzlar, Germany). The numbers of viable and aborted seeds in transgenic lines were statistically analyzed with a $\chi^2$ test. The probability that deviates from a 1:1 or 3:1 segregation ratio for viable and aborted seeds was also calculated.

We found *dme-2*/*dme-2* Col-*gl* plants can be easily obtained from *DME*/*dme-2* heterozygotes if seeds prior to desiccation were rescued on MS sucrose plates. This is consistent with the report that the *fis* endosperm cellularization defect and embryo arrest can be rescued by culturing the developing seeds in sucrose media because *fis* seeds have reduced hexose levels (1). Using this method, we generated multiple homozygous lines, and we did not detect any difference between individuals in terms of normal seed rate or visible phenotype. The adult *dme-2*/*dme-2* plants are morphologically indistinguishable from wild-type Col-*gl* plants but produce ~0.1% viable mature seeds. These *dme-2*/*dme-2* plants are not due to genetic mutation or heritable aberrant epigenetic effects that escape the requirement of DME activity during gametogenesis because their subsequent progeny are phenotypically normal and produces same level (~0.1%) of normal seeds. For complementation assays in *dme-2*/*dme-2* homozygous plants, seeds were harvested from dipped $T_0$ plants. Transgenic plants were identified by germinating normal $T_1$ seeds on MS plates containing hygromycin. T1 plants carrying a single locus of complementing transgene in *dme-2/dme-2* transgenic plants produced siliques with 50% seed abortion rate. By contrast, when transforming *dme-2/dme-2* plants, a non-complementing transgene does not produce any transgenic plant.

**Whole-Genome Bisulfite Sequencing and DNA Methylome Analysis.**

Genomic DNA were isolated from hand dissected, 7-9 DAP *dme-2* endosperm that has been complemented by *nDME*$^{CTD}$ (*dme-2/dme-2;nDME*$^{CTD}$*/nDME*$^{CTD}$). Whole genome bisulfite sequencing libraries were constructed as described before with modifications (2, 3). Approximately 20-50 ng of purified genomic DNA was spiked with 0.5ng of unmethylated cl857 *Sam7* Lambda DNA (Promega, Madison, WI) and sheared to about 300bp using Covaris M220 (Covaris Inc., Woburn, Massachusetts) under the following settings: target BP, 300; peak incident power, 75 W; duty factor, 10%; cycles per burst, 200; treatment time, 90 second; sample volume 50$\mu$l. The sheared DNA was cleaned up and recovered by 1.2x AMPure XP beads then followed by end repaired and A-tailing (NEBNext Ultra II DNA Library Prep Kit for Illumina, NEB) before ligated to the NEBNext methylated multiplex adaptors (NEBNext Multiplex Oligos for Illumina, NEB) according to the manufacturer's instructions. Adaptor-ligated DNA was cleaned up with 1x AMPure XP beads. The purified adaptor-ligated DNA was spiked with 50ng of unmethylated cl857 *Sam7* Lambda DNA and subjected to one round of sodium bisulfite conversion using the EZ DNA Methylation-Lightning Kit (Zymo Research Corporation, Irvine, CA) as outlined in the manufacturer's instruction with 80 min of conversion time. Half of the bisulfite-converted DNA molecules was PCR amplified with the following condition: 2.5 U of ExTaq DNA polymerase (Takara), 5 ul of 10 x Extaq reaction buffer, 25 $\mu$M dNTPs, 1 ul of universal and index primers (10 uM) in 50 uL reaction. The thermocyling condition was as follows: 95 °C for 2 min and then 10 cycles each of 95 °C for 30 s, 65 °C for 30 s, and 72 °C for 60 s. The enriched libraries were purified twice with 0.8x (v/v) AMPure XP beads to remove adaptor dimers. High throughput sequencing was performed by Novogene Corporation (USA). Sequencing reads from three individual transgenic lines were used in the analysis (Table S7). We used the combined reads from the three independent lines for subsequent analyses since they were highly concordant, with Pearson correlation coefficients between combined independent lines, ranging between 0.92 and 0.94 (Table S4, see also whole genome average methylation plot analysis (Fig. S4)), and so that all comparisons were confined to the same cutoff criteria and comparable sequencing depth coverage.

Sequenced reads were mapped to the TAIR10 reference genomes and DNA methylation analyses were performed as previously described (2). Fractional CG methylation in 50-bp windows across the genome was compared between *dme-2*, wild-type (GSE38935), *and nDME*$^{CTD}$ -complemented endosperm. Windows with a fractional CG methylation difference of at least 0.3 in the endosperm comparison (Fisher's exact test p-value < 0.001) were merged to generate larger differentially methylated regions (DMRs) if they occurred within 300 bp. DMRs were retained for further analysis if the fractional CG methylation across the merged DMR was 0.3 greater in *dme* endosperm than in wild-type or in *nDME*$^{CTD}$ -complemented endosperm (Fisher's exact test p-value < $10^{-10}$), and if the DMR is at least 100-bp long. The merged DMR list is in the supplemental Dataset S2. The *dme* and wild-type endosperm data used in this study were derived from crossed between *Col* (female parent) and *Ler* (male parent) (GSE38935). To avoid potential ecotype-specific methylation difference, *Ler* hyper-DMRs relative to Col-0 endosperms (GSE52814) were identified using the same criteria as described above and excluded from further analyses. The DMR Venn diagrams are made based on the number of the merged DMRs.

The *ssrp1*/+ endosperm sequencing reads (4), (GSE10500) were mapped to the TAIR10 reference genome. We compared ssrp1/+ and wild-type endosperm (GSE38935) to identify SSRP1 DMRs using the method described above. DME DMRs that intersect with SSRP1 DMRs (using BEDTools intersect function (5)) were determined as FACT-dependent DME DMRs, and the ones that do not intersect with SSRP1 DMRs were determined as FACT-independent DME DMRs. For further analysis of FACT-dependent loci, we only focus on the DME DMR windows that overlap with SSRP1 DMRs.

To investigate the distribution of DMRs along genes, all TAIR-annotated genes were aligned at the 5' end or the 3' ends. The proportion of genes with specific DMRs in each 100-bp interval is plotted (Fig. 4B).

Genomic coordinates of chromatin states were from (6), converted to 50 bp windows and aligned with *dme-2*, wild-type and $nDME^{CTD}$ methylomes. Each 50 bp windows of wild-type DME and $nDME^{CTD}$ DMRs was assigned a chromatin state accordingly.

Methylation average metaplots of Genes and TEs (Fig. S4) were performed as described previously (2).

**RNA extraction, cDNA synthesis and quantitative PCR analysis.**

Total RNA was extracted using TRIzol® Reagent (Invitrogen, Carlsbad, USA) and treated with TURBO DNase (Ambion, Austin TX, USA) according to the manufacturers' instructions. For cDNA synthesis, 5mg of total RNA was reverse-transcribed using Superscript III Reverse Transcriptase and oligo(dT) primer (Invitrogen). cDNA was treated with RNase H (Invitrogen) at 37°C for 20min and diluted tenfold with $H_2O$. For each 15-$\mu$l qPCR reaction, 1$\mu$l of diluted cDNA was used. Quantitative RT-PCR (qRT-PCR) was run on an ABI 7500 Fast Real-Time PCR System (http://www.appliedbiosystems.com) using FastStart Universal SYBR Green Master Mix (Roche, http://www.roche.com). The qRT-PCR primers are listed in Table S3. Ct values were normalized against *ACT2* (*At3g18780*) mRNA or *UBC* (*At5g25760*) mRNA. The abundance of mRNAs was expressed as relative to controls, with control values set to 1. The error bars represent the standard deviation of 4 biological replicates.

**Protein domain analysis and phylogenetic inference.**

We utilized a domain-centric computational strategy to study DME and its related proteins. Specifically, we identify DME homologs by using the iterative profile searches with PSI-BLAST (7) from the protein non-redundant (NR) database at the National Center for Biotechnology Information (NCBI). Multiple sequence alignments were built by the Promals (8) program, followed by careful manual adjustments. Consensus secondary structures were predicted using the PSIPRED(9) JPred program (10). Conserved domains were further characterized based on the comparison to available domain models from pfam (11) and sequence/structural features. The PhyML program (12) was used to determine the maximum-likelihood tree using the Jones–Taylor–Thornton (JTT) model for amino acids substitution with a discrete gamma model (four categories with gamma shape parameter: 1.096). The tree was rendered using MEGA Tree Explorer (13).

### Supplementary Figures

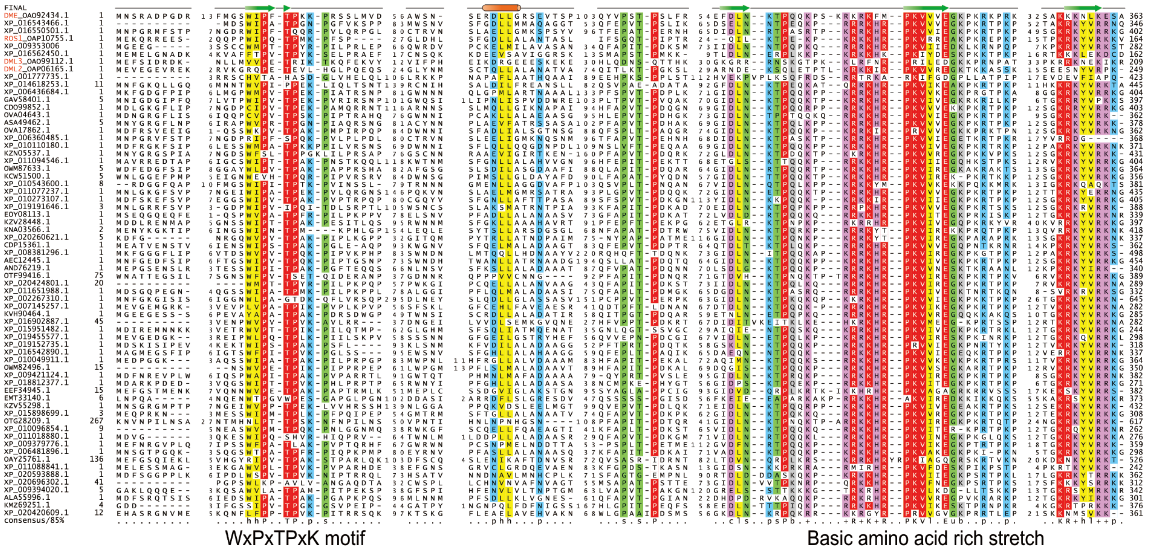


**Fig. S1.** Alignment of angiosperm DME-like proteins showing the conserved DemeN domain and basic stretch region. Bioinformatic analysis using available DME-like sequences identified a ~120-amino-acid-long conserved region at the very N-terminus of DME-like proteins in angiosperms. This sequence is characterized by a highly conserved WxPxTPxK motif that might function in protein-protein interactions. Further toward the C-terminus is a stretch of basic amino acids that serves as a nuclear localization signal. This sequence consists of direct repeats reminiscent of the AT-hook motifs that may bind DNA. Numbers flanking each sequence represent amino acid residue positions. Numbers between alignment blocks reflect residue gaps not shown in the alignment.

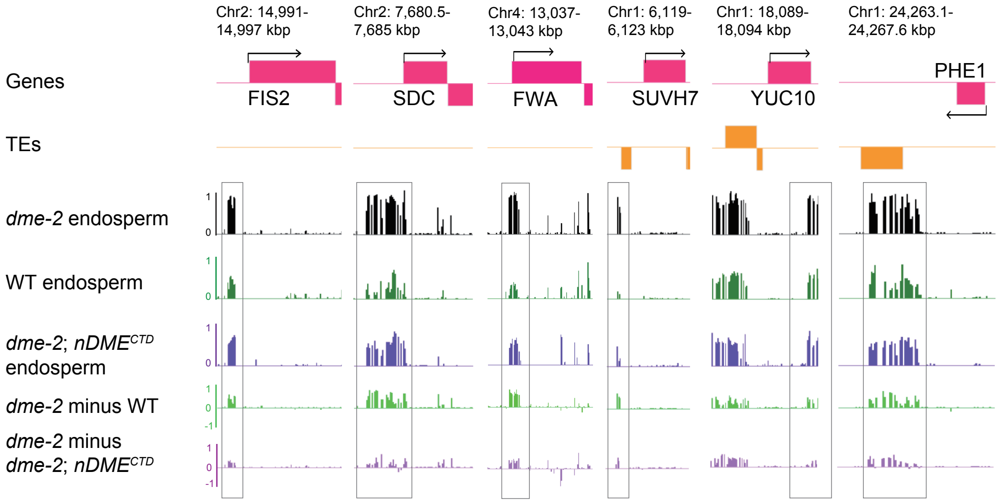


**Fig. S2** Genome browser snapshots of CG DNA methylation at selected imprinted gene loci. Top two tracks are coding genes (magenta) and TEs (orange) with Tair10 chromosome coordinates. The bottom five tracks represent fractional CG methylation levels for different genotypes: black trace, *dme-2* endosperm; dark green trace, WT endosperm; dark blue trace, *nDME*$^{CTD}$-complemented endosperm; light green trace, WT endosperm subtracted from *dme-2* mutant endosperm; light purple trace, *nDME*$^{CTD}$-complemented endosperm subtracted from *dme-2* endosperm. *dme*-induced CG hypermethylation at selected maternally expressed (*FIS2* and *SDC*) and paternally expressed (*SUVH7*, *YUC10, and PHE1*) imprinted genes is rescued in *nDME*$^{CTD}$-complemented endosperm.

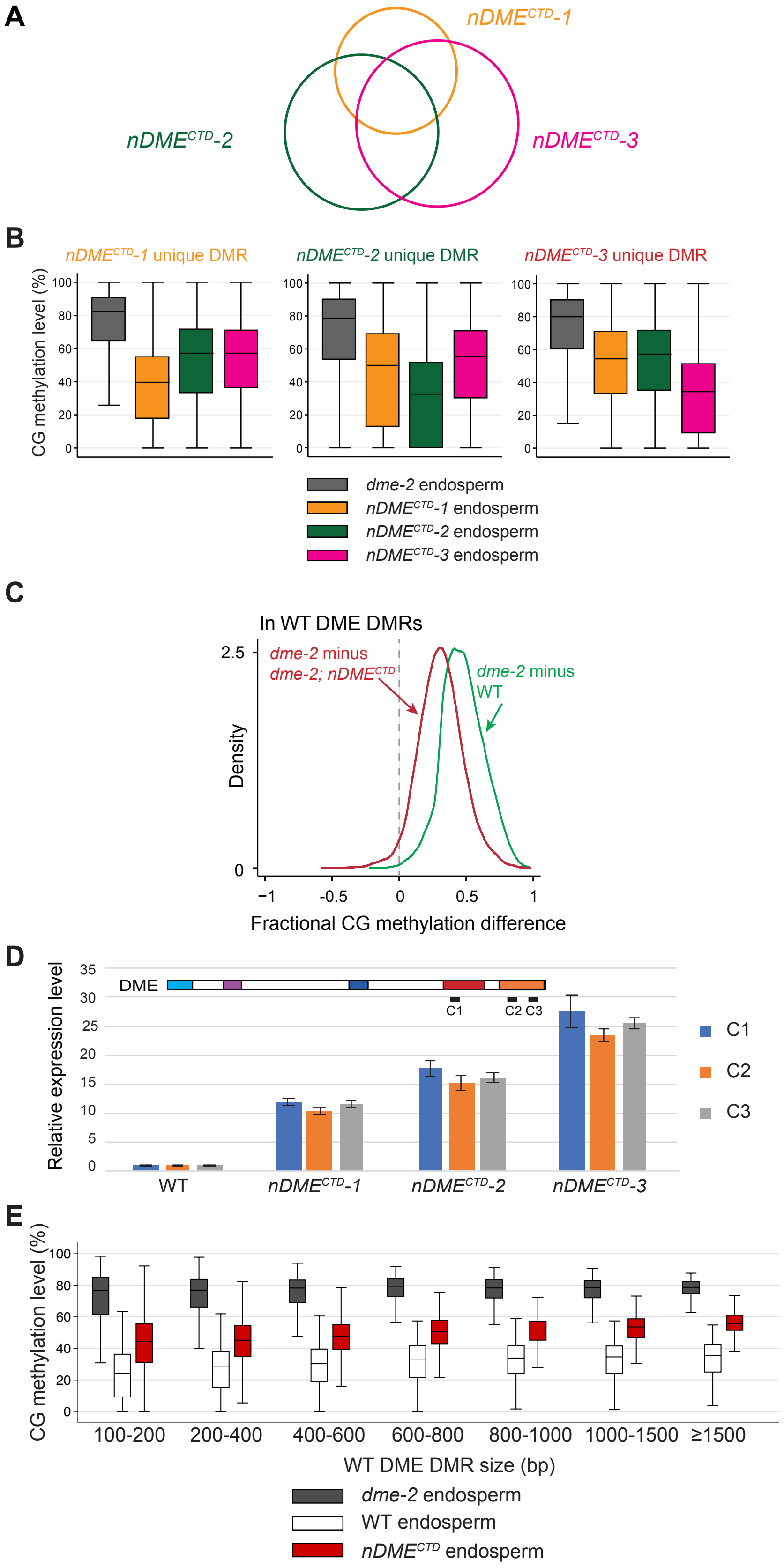
A
nDME^CTD-1
nDME^CTD-2
nDME^CTD-3
B
nDME^CTD-1 unique DMR
nDME^CTD-2 unique DMR
nDME^CTD-3 unique DMR
CG methylation level (%)
dme-2 endosperm
nDME^CTD-1 endosperm
nDME^CTD-2 endosperm
nDME^CTD-3 endosperm
C
In WT DME DMRs
dme-2 minus dme-2; nDME^CTD
dme-2 minus WT
Density
Fractional CG methylation difference
D
Relative expression level
DME
C1 C2 C3
WT
nDME^CTD-1
nDME^CTD-2
nDME^CTD-3
E
CG methylation level (%)
100-200
200-400
400-600
600-800
800-1000
1000-1500
≥1500
WT DME DMR size (bp)
dme-2 endosperm
WT endosperm
nDME^CTD endosperm

**Fig. S3.** DNA methylomes of three independent *nDME*$^{CTD}$-complemented endosperm replicates. (A) Venn diagram showing partial overlap of DMRs of each *nDME*$^{CTD}$-complemented endosperm relative to *dme-2* mutant endosperm (*nDME*$^{CTD}$*-1* to *nDME*$^{CTD}$-3). (B) Boxplot of CG methylation levels among canonical DME target sites in *dme-2* mutant (grey), wild-type (white), or in combined *nDME*$^{CTD}$- complemented (red) endosperm. (C) Kernel density plot of CG methylation differences between *dme* and *nDME*$^{CTD}$-complemented endosperm, for loci demethylated by DME. Green trace shows these DME target sites are hypermethylated in *dme* endosperm. Red trace shows these same DME target sites are partially demethylated by nDME$^{CTD}$. (D) qRT-PCR of *nDME*$^{CTD}$ expression levels in the three independent nDME$^{CTD}$-complemented lines used in this study compared to that of the endogenous DME. (E) Boxplot of CG methylation levels within canonical DME target sites grouped by DMR length, in *dme-2* mutant (black), wild-type (white), or *nDME*$^{CTD}$-complemented (red) endosperm.

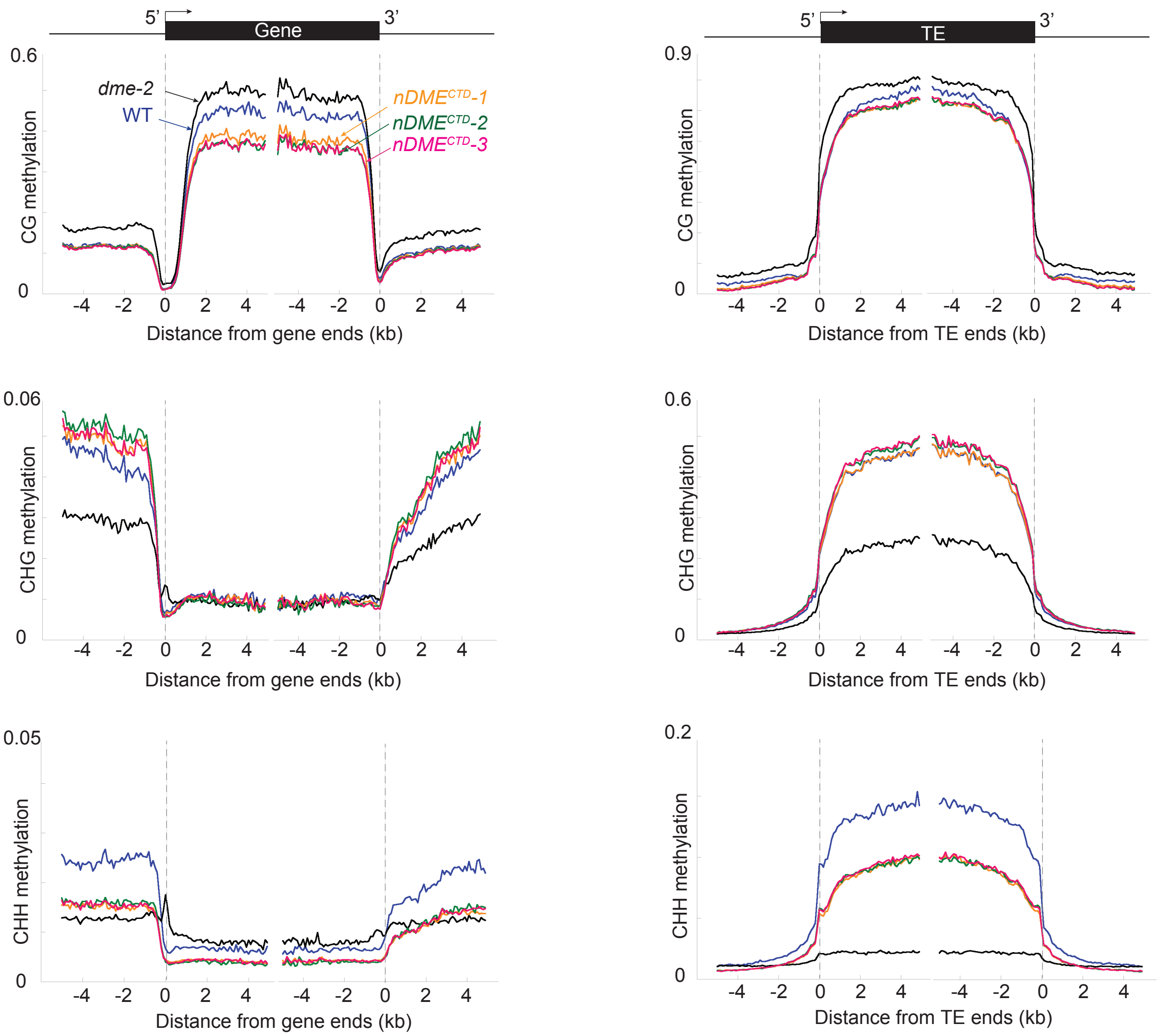


**Fig. S4.** Whole genome CG, CHG, CHH methylation average plots of three independent nDME$^{CTD}$-complemented lines, *dme-2* mutant, and wild-type endosperm. Average CG, CHG, CHH methylation in Genes (left panels) or TEs (right panels) of endosperm methylation data used in this study. Wildtype (WT) and *dme-2* endosperm data were from (2). Three nDME$^{CTD}$-complemented endosperm methylation profiles were as indicated. Endosperm were dissected from complemented lines from bent-cotyledon stage seeds.

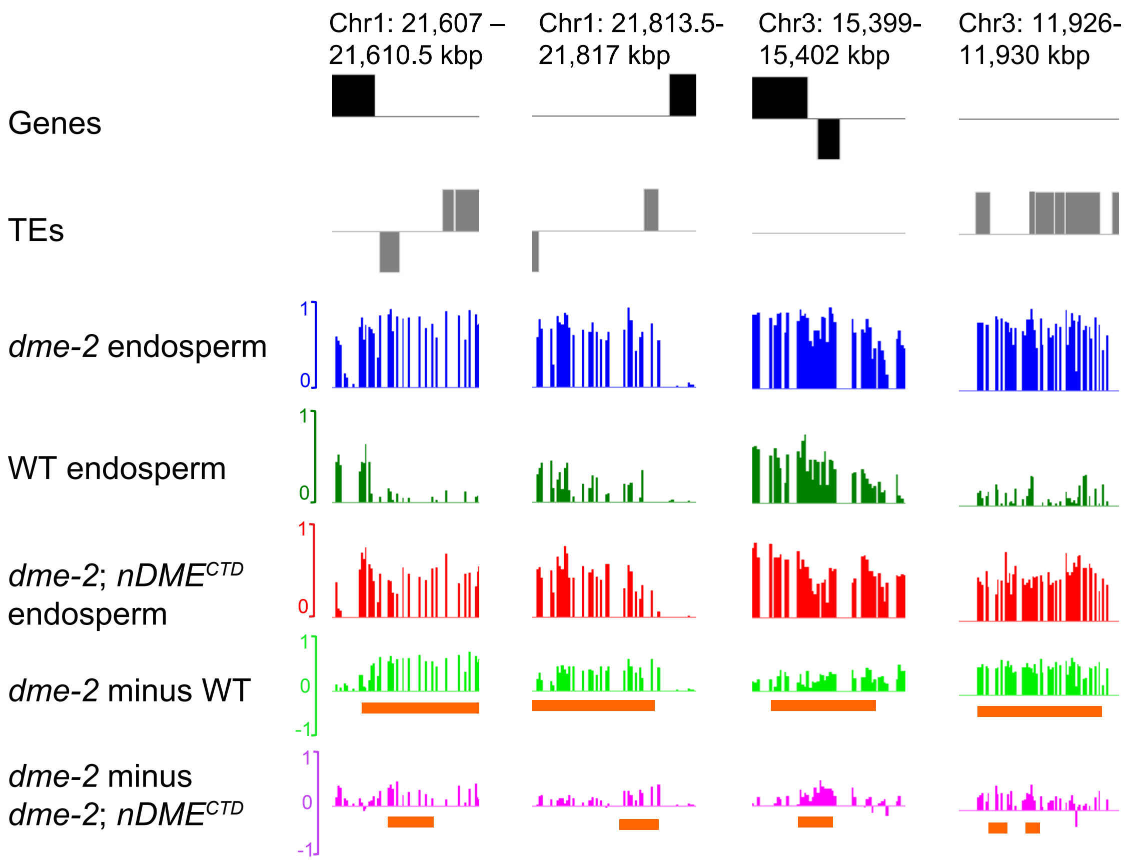


**Fig. S5.** Genome Browser snapshots of long wild-type DME DMRs. Tracks are as labeled. The DMR regions are indicated as horizontal bars (bottom two tracks) according to their length in each sample.

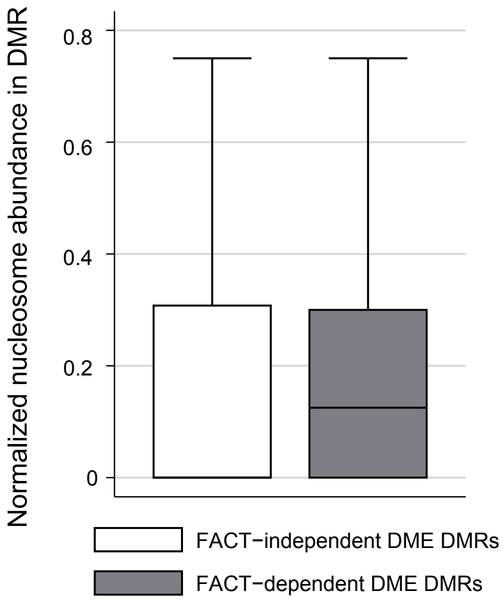


**Fig. S6.** Boxplot of relative nucleosome abundance in FACT-independent or -dependent DME DMRs, using genome coordinates of the most well-positioned nucleosomes (14) that overlap with DME DMRs. Number of nucleosomes per 100-bp DMR is plotted

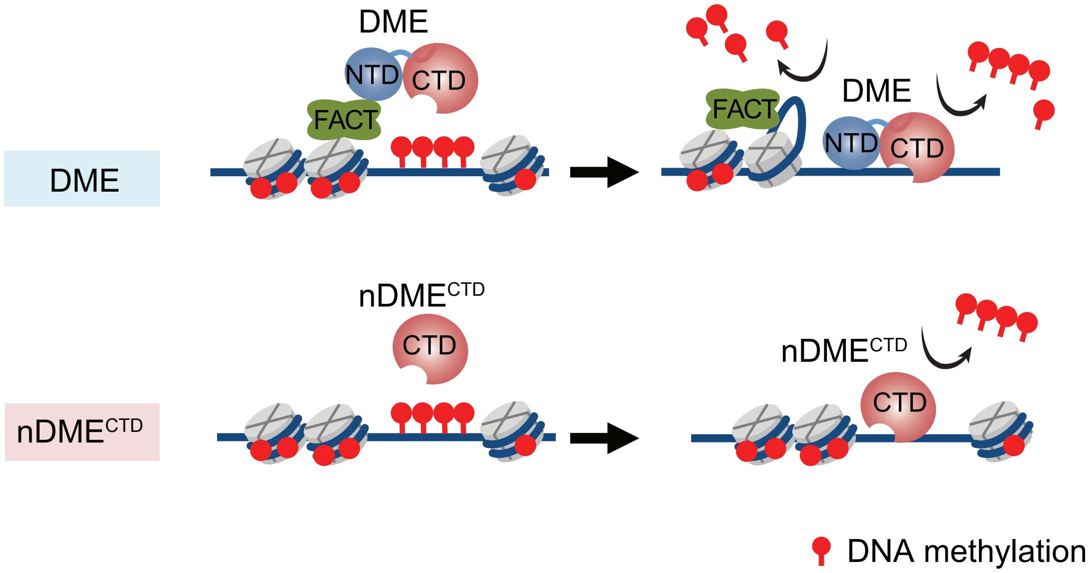


**Fig. S7.** A bipartite model for DME-mediated active demethylation. We propose a bipartite organization of DME where the targeting and recruitment information is contained within the C-terminal catalytic core. In the absence of the N-terminal domain, processing of heterochromatin demethylation is significantly impeded, presumably by chromatin structure. The N-terminal domain is required to recruit FACT complex for the heterochromatic targets to overcome nucleosomal obstacles. The N-terminal conserved domains were acquired late during land plant evolution and are restricted to the angiosperm linage, suggesting a mode of demethylation regulation specific to angiosperm genomes.

### Supplementary Tables

### Table S1. Seed abortion ratio of *dme-2* complemented by *nDME*$^{CTD}$

| Genotype | No. of total seeds* | No. of viable seeds | Proportion of viable seeds (%) | *p* for 1:1† | *p* for 3:1† |
|---|---|---|---|---|---|
| *DME/dme-2; pDME:nDME$^{CTD}$-2* | 347 | 287 | 83 | 3.7E-34 | 9.1E-04 |
| *DME/dme-2; pDME:nDME$^{CTD}$-1* | 444 | 372 | 84 | 5.4E-46 | 1.9E-05 |
| *DME/dme-2; pDME:nDME$^{CTD}$-3* | 171 | 119 | 70 | 3.0E-07 | 0.10 |
| *DME/dme-2; pDME:nDME$^{CTD}$-4* | 124 | 80 | 65 | 1.2E-03 | 0.01 |
| *DME/dme-2; pDME:nDME$^{CTD}$-45* | 282 | 213 | 76 | 9.9E-18 | 0.84 |
| *DME/dme-2; pDME:nDME$^{CTD}$-5* | 363 | 221 | 61 | 3.4E-05 | 5.2E-10 |
| *dme-2/dme-2; pDME:nDME$^{CTD}$-6* | 221 | 166 | 75 | 8.2E-14 | 0.97 |
| *dme-2/dme-2; pDME:nDME$^{CTD}$-8* | 284 | 206 | 73 | 3.1E-14 | 0.34 |
| *dme-2/dme-2; pDME:nDME$^{CTD}$-4* | 221 | 155 | 70 | 2.1E-09 | 0.09 |
| *dme-2/dme-2; pDME:nDME$^{CTD}$-11* | 413 | 269 | 65 | 7.7E-10 | 3.6E-06 |
| *dme-2/dme-2; pDME:nDME$^{CTD}$-3* | 351 | 206 | 59 | 1.1E-03 | 1.7E-12 |
| *dme-2/dme-2; pDME:nDME$^{CTD}$-16* | 229 | 131 | 57 | 0.03 | 5.0E-10 |
| *dme-2/dme-2; pDME:nDME$^{CTD}$15* | 413 | 208 | 50 | 0.88 | 6.4E-31 |
| *dme-2/dme-2; pDME:nDME$^{CTD}$-1* | 298 | 150 | 50 | 0.91 | 8.1E-23 |
| *dme-2/dme-2; pDME:nDME$^{CTD}$-10* | 300 | 151 | 50 | 0.91 | 5.8E-23 |
| *dme-2/dme-2; pDME:nDME$^{CTD}$-13* | 387 | 186 | 48 | 0.44 | 1.9E-34 |
| *dme-2/dme-2; pDME:nDME$^{CTD}$-12* | 223 | 83 | 37 | 1.3E-04 | 8.3E-39 |

* Total number of viable and aborted seed counted.

† Probability that the deviation from the indicated segregation ratio of viable: aborted seeds is due to chance.

### Table S2. Reduced paternal *dme* allele transmission in the Col-*gl* ecotype

| Genotype | Ecotype | *DME/dme-2* * | *DME/DME* * | *dme-2* (%) § | *p* † |
|---|---|---|---|---|---|
| *DME/dme-2* self-pollination, plant 1 | Col-*gl* | 78 | 190 | 29 | 7.8E-12 |
| *DME/dme-2* self-pollination, plant 2 | Col-*gl* | 62 | 172 | 26 | 6.4E-13 |
| *DME/dme-2* self-pollination, plant 3 | Col-*gl* | 72 | 145 | 33 | 7.2E-7 |
| *DME/dme-2* self-pollination, plant 4 | Col-*gl* | 51 | 201 | 20 | 3.4E-21 |

* Number of F1 seed with indicated genotype.

§ *dme-2* allele transmission rate.

† Probability that that the deviation from the indicated segregation ration (1:1 inheritance of paternal genome with or without *dme-2* in the F1 generation) is due to chance.

### Table S3. Complementation of *dme* allele pollen transmission by nDME$^{CTD}$

| Maternal parent | Paternal parent | Paternal parent normal seed (%) | *DME/dme-2* * | *DME/dme-2*; *T* * | *T* (%) § | *p* † |
|---|---|---|---|---|---|---|
| Col-0 | *dme-2/dme-2*; *nDME$^{CTD}$* Line 10 | 50 | 32 | 62 | 66 | 2.0E-3 |
| Col-0 | *dme-2/dme-2*; *nDME$^{CTD}$* Line 11 | 65 | 3 | 50 | 94.3 | 1.1E-10 |
| Col-0 | *dme-2/dme-2*; *nDME$^{CTD}$* Line 12 | 37 | 8 | 34 | 81 | 6.0E-5 |
| Col-0 | *dme-2/dme-2*; *nDME$^{CTD}$* Line 15 | 50 | 9 | 44 | 83 | 1.5E-6 |

* Number of F1 seed with indicated genotype. *T*, *nDME$^{CTD}$* transgene.

§ *T*, *nDME$^{CTD}$* transgene transmission rate.

† Probability that that the deviation from the indicated segregation ration (1:1 inheritance of paternal genome with or without *nDME$^{CTD}$* transgene in the F1 generation) is due to chance.

### Table S4. Whole genome endosperm BS-seq data correlation between three independent complementation lines

| Pearson correlation coefficient | *nDME$^{CTD}$-1* endosperm CG | *nDME$^{CTD}$-2* endosperm CG | *nDME$^{CTD}$-3* endosperm CG |
|---|---|---|---|
| *nDME$^{CTD}$-1* endosperm CG | 1 | | |
| *nDME$^{CTD}$-2* endosperm CG | 0.9423 | 1 | |
| *nDME$^{CTD}$-3* endosperm CG | 0.9317 | 0.9331 | 1 |

### Table S5. Whole genome endosperm BS-seq data correlation between the three data sets used in this study excluding canonical DME target sites

| Pearson correlation coefficient | *dme-2* endosperm CG | WT endosperm CG | *nDME$^{CTD}$* endosperm CG |
|---|---|---|---|
| *dme-2* endosperm CG | 1 | | |
| WT endosperm CG | 0.9348 | 1 | |
| *nDME$^{CTD}$* endosperm CG | 0.9434 | 0.9244 | 1 |

### Table S6. Gene enrichment analysis of *nDME*$^{CTD}$-unique target genes

| GO Function | Total gene | *nDME*$^{CTD}$-unique target gene | Expected | Fold Enrichment | FDR (<0.05) |
|---|---|---|---|---|---|
| **Biological Process** | | | | | |
| gluconeogenesis (GO:0006094) | 17 | 6 | 1 | 4.38 | 2.62E-02 |
| regulation of gene expression, epigenetic (GO:0040029) | 45 | 12 | 4 | 3.31 | 4.78E-03 |
| DNA replication (GO:0006260) | 129 | 25 | 10 | 2.41 | 1.89E-03 |
| tRNA metabolic process (GO:0006399) | 143 | 26 | 12 | 2.26 | 3.44E-03 |
| RNA splicing, via transesterification reactions (GO:0000375) | 202 | 36 | 16 | 2.21 | 4.09E-04 |
| nuclear transport (GO:0051169) | 119 | 21 | 10 | 2.19 | 9.86E-03 |
| phospholipid metabolic process (GO:0006644) | 110 | 19 | 9 | 2.14 | 1.89E-02 |
| mRNA processing (GO:0006397) | 310 | 52 | 25 | 2.08 | 7.04E-05 |
| mRNA splicing, via spliceosome (GO:0000398) | 224 | 37 | 18 | 2.05 | 1.39E-03 |
| DNA repair (GO:0006281) | 189 | 31 | 15 | 2.04 | 3.78E-03 |
| DNA metabolic process (GO:0006259) | 351 | 57 | 28 | 2.02 | 4.64E-05 |
| protein transport (GO:0015031) | 600 | 97 | 48 | 2.01 | 4.30E-08 |
| intracellular protein transport (GO:0006886) | 568 | 90 | 46 | 1.97 | 4.52E-07 |
| phosphate-containing compound metabolic process (GO:0006796) | 1688 | 259 | 136 | 1.9 | 4.34E-19 |
| vesicle-mediated transport (GO:0016192) | 520 | 78 | 42 | 1.86 | 1.54E-05 |
| transcription from RNA polymerase II promoter (GO:0006366) | 427 | 63 | 34 | 1.83 | 2.23E-04 |
| cellular amino acid biosynthetic process (GO:0008652) | 185 | 27 | 15 | 1.81 | 2.66E-02 |
| nucleobase-containing compound metabolic process (GO:0006139) | 2727 | 397 | 220 | 1.81 | 4.23E-26 |
| protein localization (GO:0008104) | 474 | 68 | 38 | 1.78 | 2.06E-04 |
| cell cycle (GO:0007049) | 532 | 76 | 43 | 1.77 | 1.04E-04 |
| regulation of nucleobase-containing compound metabolic process (GO:0019219) | 409 | 58 | 33 | 1.76 | 9.26E-04 |
| RNA metabolic process (GO:0016070) | 1603 | 227 | 129 | 1.76 | 3.94E-13 |
| catabolic process (GO:0009056) | 1290 | 176 | 104 | 1.69 | 5.58E-09 |
| transcription, DNA-dependent (GO:0006351) | 753 | 99 | 61 | 1.63 | 1.05E-04 |
| transport (GO:0006810) | 1441 | 187 | 116 | 1.61 | 4.46E-08 |
| cellular protein modification process (GO:0006464) | 783 | 99 | 63 | 1.57 | 4.09E-04 |
| cellular component organization (GO:0016043) | 1359 | 169 | 110 | 1.54 | 2.87E-06 |
| localization (GO:0051179) | 1580 | 194 | 127 | 1.52 | 8.15E-07 |
| nitrogen compound metabolic process (GO:0006807) | 2924 | 359 | 236 | 1.52 | 5.08E-13 |
| cellular component organization or biogenesis (GO:0071840) | 1640 | 198 | 132 | 1.5 | 1.62E-06 |
| primary metabolic process (GO:0044238) | 5340 | 631 | 430 | 1.47 | 3.18E-21 |
| metabolic process (GO:0008152) | 6924 | 811 | 558 | 1.45 | 9.32E-28 |
| organelle organization (GO:0006996) | 938 | 108 | 76 | 1.43 | 3.71E-03 |
| cellular process (GO:0009987) | 6866 | 780 | 553 | 1.41 | 4.33E-23 |
| regulation of biological process (GO:0050789) | 1302 | 144 | 105 | 1.37 | 2.59E-03 |
| biological regulation (GO:0065007) | 1526 | 164 | 123 | 1.33 | 3.16E-03 |
| response to stimulus (GO:0050896) | 1690 | 178 | 136 | 1.31 | 3.73E-03 |
| protein metabolic process (GO:0019538) | 1751 | 181 | 141 | 1.28 | 6.73E-03 |
| biosynthetic process (GO:0009058) | 2090 | 206 | 168 | 1.22 | 2.36E-02 |

**Table S6. Continued.**

| | | | | | |
|---|---|---|---|---|---|
| **Molecular Function** | | | | | |
| lipid transporter activity (GO:0005319) | 20 | 7 | 1.61 | 4.34 | 1.95E-02 |
| enzyme activator activity (GO:0008047) | 46 | 14 | 3.71 | 3.78 | 1.26E-03 |
| nucleotidyltransferase activity (GO:0016779) | 128 | 30 | 10.31 | 2.91 | 3.60E-05 |
| DNA-directed RNA polymerase activity (GO:0003899) | 60 | 14 | 4.83 | 2.9 | 8.04E-03 |
| helicase activity (GO:0004386) | 87 | 19 | 7.01 | 2.71 | 2.72E-03 |
| DNA helicase activity (GO:0003678) | 52 | 11 | 4.19 | 2.63 | 3.38E-02 |
| microtubule motor activity (GO:0003777) | 75 | 15 | 6.04 | 2.48 | 2.21E-02 |
| anion channel activity (GO:0005253) | 77 | 14 | 6.2 | 2.26 | 4.16E-02 |
| translation regulator activity (GO:0045182) | 105 | 19 | 8.46 | 2.25 | 1.93E-02 |
| motor activity (GO:0003774) | 94 | 17 | 7.57 | 2.24 | 2.98E-02 |
| hydrogen ion transmembrane transporter activity (GO:0015078) | 102 | 18 | 8.22 | 2.19 | 2.55E-02 |
| pyrophosphatase activity (GO:0016462) | 609 | 107 | 49.07 | 2.18 | 2.15E-10 |
| GTPase activity (GO:0003924) | 194 | 34 | 15.63 | 2.18 | 1.15E-03 |
| ion channel activity (GO:0005216) | 93 | 16 | 7.49 | 2.14 | 4.40E-02 |
| mRNA binding (GO:0003729) | 177 | 28 | 14.26 | 1.96 | 1.33E-02 |
| nucleotide binding (GO:0000166) | 141 | 22 | 11.36 | 1.94 | 3.49E-02 |
| kinase activity (GO:0016301) | 724 | 111 | 58.34 | 1.9 | 7.19E-08 |
| receptor activity (GO:0004872) | 361 | 54 | 29.09 | 1.86 | 8.57E-04 |
| transmembrane receptor protein serine/threonine kinase activity (GO:0004675) | 334 | 49 | 26.91 | 1.82 | 2.47E-03 |
| signal transducer activity (GO:0004871) | 305 | 44 | 24.58 | 1.79 | 5.60E-03 |
| protein kinase activity (GO:0004672) | 508 | 71 | 40.93 | 1.73 | 5.69E-04 |
| transferase activity (GO:0016740) | 1902 | 252 | 153.26 | 1.64 | 1.68E-11 |
| RNA binding (GO:0003723) | 739 | 86 | 59.55 | 1.44 | 1.16E-02 |
| hydrolase activity (GO:0016787) | 2086 | 238 | 168.08 | 1.42 | 8.75E-06 |
| binding (GO:0005488) | 3418 | 383 | 275.41 | 1.39 | 7.64E-09 |
| nucleic acid binding (GO:0003676) | 1759 | 195 | 141.73 | 1.38 | 4.22E-04 |
| catalytic activity (GO:0003824) | 5737 | 635 | 462.26 | 1.37 | 7.30E-15 |
| protein binding (GO:0005515) | 1594 | 175 | 128.44 | 1.36 | 1.17E-03 |
| transporter activity (GO:0005215) | 996 | 108 | 80.25 | 1.35 | 2.50E-02 |

### Table S7. Primer list

| | |
|---|---|
| Primers used for plasmid construction: | |
| VeDME | AGTGGAGACGGACGTCCTGACGCCCTCAAAAATGTCTTCTTAGGATCACAAAATC |
| P3R | CACCATTGTTAACACACTTGATGAATCACTCCCCCTTC |
| lnAGBF | GCTGCCGCAGCGGCTGCCTACAAAGGAGATGGTGCACTTGTT |
| CTDVeR | AGGACTCTAGGGACTAGTCCCGGGTTTAGGTTTTGTTGTTCTTCAATTTGCTCGCAG |
| ATGF | TTCATCAAGTGTGTTAACAATGGTGGCTGCCGCAGCGGCTGCCTACAAAG |
| ATGR | CTTTGTAGGCAGCCGCTGCGGCAGCCACCATTGTTAACACACTTGATGAA |
| S40F | TTCATCAAGTGTGTTAACAATGGTGCCAAAGAAGAAGAGAAAGGTC |
| S40R | CTTTGTAGGCAGCCGCTGCGGCAGCGACCTTTCTCTTCTTCTTTGG |
| Primers for qRT-PCR analysis: | |
| qFIS2F | TCGATTGGTGGTGGAGAATG |
| qFIS2R | AGTTACTGAGGAGGATGGTAGT |
| qFWAF | CGCCTTCTTCTTCCTCTAATCC |
| qFWAR | AGTCCACTTCTCCCAATGTAATC |
| ACT2F | GACCTTTAACTCTCCCGCTATG |
| ACT2R | GAGACACACCATCACCAGAAT |
| uUBCF | CTGCGACTCAGGGAATCTTCTAA |
| uUBCR | TTGTGCCATTGAATTGAACCC |
| qDMEc4F | GTGGTTGATCCGCTCAGTAA |
| qDMEc4R | CGTCCCTTTCATCTCTCGTAAA |
| qDMEc5F | GAGGAGAGGAGCTTAACAAGTG |
| qDMEc5R | TCTCCAACGGAAGAGGTAGT |
| qDMEc6F | CATCGTCTCCTTGATGGTATGG |
| qDMEc6R | CTTTCCCTCCACACTTCTGTT |

**Table S8. Average genomic coverage and DNA methylation for the samples in this study**

| Sample | Raw reads | Aligned bases | Average coverage | Chloroplast CHH methylation (%) |
|---|---|---|---|---|
| *dme-2/dme-2; nDME$^{CTD}$-1/nDME$^{CTD}$-1* | 34935210 | 3746188650 | 15 | 0.3 |
| *dme-2/dme-2; nDME$^{CTD}$-2/nDME$^{CTD}$-2* | 38167678 | 4239339600 | 17 | 0.3 |
| *dme-2/dme-2; nDME$^{CTD}$-3/nDME$^{CTD}$-3* | 37046004 | 3999866100 | 16 | 0.3 |

Chloroplast CHH methylation is an estimate of the bisulfite non-conversion rate in the sequenced samples.
